# ICON: An isoform-aware hierarchical random forest model for cell type classification

**DOI:** 10.64898/2026.04.28.721306

**Authors:** Hettiarachchige Wijewardena, Siyuan Wu, Ulf Schmitz

## Abstract

Single-cell RNA sequencing (scRNA-seq) has transformed our ability to resolve cellular heterogeneity across complex biological systems. However, conventional short-read scRNA-seq is inherently limited in its inability to capture full-length transcripts. Isoform profiles, arising from alternative splicing, provide a deeper layer of resolution, enabling finer discrimination of cellular subtypes and dynamic states, particularly in heterogenous tissues. Long-read RNA sequencing technologies enable accurate transcript-level profiling and more comprehensive characterisation of isoform diversity. Despite these advances, existing cell type annotation methods remain largely tailored to gene-level data, thereby limiting fidelity and leaving isoform-level information an untapped reservoir of biological insight.

Here, we present a hierarchical random forest (HRF) framework, ICON, for isoform-aware cell classification in scRNA-seq data. By jointly modelling gene- and isoform-level expression the framework captures both abundance and useage patterns, enabling classification beyond gene-level resolution. A two-stage strategy first assigns cell identities using highly variable gene and isoform features, followed by targeted reclassification of ambiguous cells based on relative isoform and gene usage, thereby resolving conflicts that arise from transcriptional heterogeneity.

Importantly, ICON provides interpretable outputs by identifying key genes and isoforms that drive cell type discrimination, linking classification to underlying regulatory mechanisms. Benchmarking on long-read scRNA-seq datasets demonstrates consistent improvements over conventional gene-based approaches. With increasing adoption of long-read sequencing, our framework provides a robust, interpretable foundation for isoform-aware cell type annotation, improving resolution and insight.

## 1. Introduction

Cells are the fundamental units of life, and their identity and function are determined by the genes and proteins they express at a given time. Variations in gene expression are closely associated with phenotypic differences and are often implicated in disease. Beyond gene-level regulation, alternative splicing generates multiple transcript isoforms from a single gene, adding an additional layer of complexity to cellular function. While orchestrated isoform expression is key for controlling critical biological processes, dysregulation of isoform usage has been linked to various diseases, including cancer [1].

Characterising the transcriptomic profiles of distinct cell types is therefore critical for understanding biological systems, identifying early disease markers, and discovering therapeutic targets. Accurate cell type identification underpins emerging applications such as somatic cell reprogramming [2], directed differentiation of pluripotent stem cells [3], and cell-based therapies, including engineered T-cell approaches [4].

scRNA-seq has enabled high-throughput profiling of gene expression at the resolution of individual cells, allowing the exploration of cellular heterogeneity within complex tissues. Current single-cell technologies, including droplet-, plate-, and microwell-based platforms, combined with advances in sequencing technologies, facilitate transcriptomic analysis at gene, isoform, and even spatial levels [5–7]. In particular, isoform-level analysis provides enhanced resolution for distinguishing closely related cell subtypes and dynamic cellular states, which is especially valuable in heterogeneous biological systems [8,9].

However, short-read scRNA-seq technologies are limited in their ability to reconstruct full-length transcripts, restricting accurate isoform-level characterisation [10]. Recent developments in long-read sequencing technologies have addressed this limitation by enabling direct sequencing of full-length transcripts [11,12]. These approaches provide improved resolution of transcript structures, facilitating the identification of novel isoforms, fusion transcripts, and other complex RNA processing events, thereby enhancing our understanding of transcriptomic diversity at the single-cell level [13,14].

Despite these technological advances, current computational approaches for cell type annotation in single-cell studies remain largely reliant on gene-level expression and clustering-based methods [15]. In most workflows, clustering is used to define cell populations, followed by manual or semi-automated annotation based on known marker genes [16,17]. Widely used frameworks such as *Seurat* first perform unsupervised clustering based on gene-expression similarity and cell populations are subsequently annotated using known marker genes or reference datasets[18]. More recently, methods such as *CHOIR* extend clustering by using HRF classifiers with permutation testing to iteratively split and merge clusters, aiming to identify statistically distinct cell populations while reducing over clustering [19]. Although powerful for population discovery, both approaches remain fundamentally reliant on gene-level representations rather than transcript isoforms.

Cell type annotation tools such as *ACTINN [20]* and *CaSTLe [21]* automate label transfer from previously annotated reference datasets. *ACTINN* uses a neural network trained on labelled gene-expression matrices to predict cell identities in new datasets, whereas *CaSTLe* applies an XGBoost-based transfer-learning framework that learns informative gene features from reference data and assigns labels to query cells. While these methods reduce the need for manual annotation, they were primarily developed for short-read, gene-level scRNA-seq data and do not fully exploit isoform-resolved expression matrices. Consequently, their performance may be limited for long-read sequencing datasets, where transcript diversity and alternative splicing provide biologically meaningful signals beyond conventional gene-level summaries [11,15]. These limitations highlight the need for dedicated isoform-aware cell type annotation approaches capable of leveraging long-read transcriptomic information.

To address these limitations, we developed a HRF framework, ICON (Isoform-aware Cell classificatiON), for cell type annotation that integrates both gene- and isoform-level information. In addition to improving classification accuracy, ICON emphasises interpretability by identifying the most informative genes and isoforms contributing to each prediction. We benchmarked the model against existing gene-based annotation tools using both gene-level and isoform-level datasets derived from long-read sequencing. Our results demonstrate that while gene-based methods perform well on gene-level data, their performance declines substantially on isoform-level matrices. In contrast, the proposed approach consistently achieves higher accuracy across both data types, highlighting the importance of incorporating isoform-level information for robust cell type annotation.

## 2. Methods

### 2.1 Dataset preprocessing

Four long-read scRNA-seq datasets were utilised for tool development and evaluation (Supplementary Table 1). For each dataset, matched gene-level and isoform-level count matrices generated from the same cells and sequencing runs were analysed on identical biological samples to enable direct comparison between gene-based and isoform-based annotation performance. Count matrices were processed in R using *Seurat* (v5.1.0). *Seurat* objects were created after filtering features detected in at least 10 cells, while retaining all cells. Data were normalised using the *LogNormalize* method, and the top 500 highly variable genes or isoforms were identified using the variance stabilising transformation (VST) method. The data were subsequently scaled, and principal component analysis (PCA) was performed for dimensionality reduction, with elbow plots used to guide principal component selection. Cell type annotations were incorporated by matching external barcode metadata to the corresponding cells. The resulting normalised, scaled, and annotated datasets were used for downstream Machine Learning (ML) –based cell type classification and benchmarking.

To preserve cell alignment and maintain consistency, no cells were removed during preprocessing. Although low-abundance features were excluded, all cells were retained to maximise data use and avoid unintended loss of rare or low-count cell types.

A comparative overview of isoform and gene expression profiles across scRNA-seq datasets, including cell type distribution (bar charts), PCA-based dimensionality reduction, and elbow plots for optimal component selection, is provided in the Supplementary Figures 1-5.

### 2.2 Evaluation metrics

Various evaluation metrics were employed to assess model performance at different stages of development. These metrics include accuracy, macro F1 score, and weighted F1 score. Accuracy was used to reflect overall classification performance aggregated across all cell types, whereas precision and recall provide cell-type–specific performance measures, which were used to compute F1 scores. The evaluation metrics are defined as follows:

Let *C* denote the total number of cell types. For each cell type *i*, let TP_*i*_, FP_*i*_ and FN_*i*_ represent the number of true positives, false positives, and false negatives, respectively.

#### Accuracy

Accuracy measures the proportion of correctly classified instances among all instances:

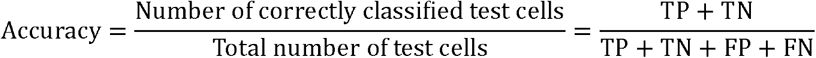

#### Precision and Recall

For cell type *i*, precision and recall are defined as:

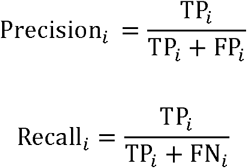

#### F1 Score

The F1 score for cell type *i* is the harmonic mean of precision and recall:

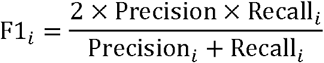

#### Macro F1 Score

The macro F1 score is calculated as the unweighted average of the F1 scores across all cell types:

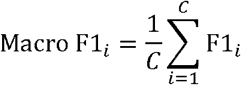

Here, *C* represents the total number of cell types (classes) in the dataset, and F1*i* is the F1 score for cell type *i*. This metric treats all cell types equally, regardless of the number of cells in each type.

#### Weighted F1 Score

The weighted F1 score accounts for class (cell count in each cell type) imbalance by weighting each class-specific F1 score by the number of true samples in that class. Let *n*_*i*_ be the number of true cells in class (cell type) *i* and *N* represents the total number of true cells across all cell types:

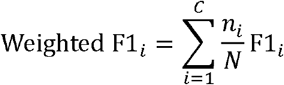

### 2.3 Random forest model parameter optimisation

RF hyperparameters, including the number of trees and number of input features, were optimised using the LongBench PacBio and Nanopore datasets. Model performance was assessed using accuracy, macro F1 score, and weighted F1 score. Full optimisation procedures are provided in the Supplementary Materials.

### 2.4 Relative isoform usage and relative gene usage

To support the hierarchical feature engineering framework, we developed two feature representations: relative isoform usage (RIU) and relative gene usage (RGU), which quantify the proportional expression of each isoform or gene within a given cell type.

Let *i* denote the specific isoform of interest, *j* an index variable used for summation over all isoforms, and *J* the total number of isoforms considered within a given cell type *c*. The relative isoform usage for isoform *i* in cell type *c* was defined as:

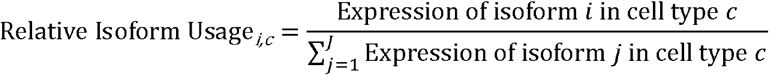

The relative gene usage was calculated in a similar manner. Genes and isoforms were ranked according to RIU or RGU values, and top-ranked features were selected for downstream modelling. More detailed methodology is provided in the Supplementary materials.

### 2.5 Model development pipeline

#### Dataset partitioning strategy for hierarchical random forest model

Model performance was evaluated using 5-fold cross-validation. In this approach, each dataset was partitioned into five approximately equal, non-overlapping folds while preserving the distribution of cell types across folds. During each iteration, four folds were used for training, and the remaining fold was used for testing. This process was repeated five times for each dataset so that each fold served once as the test set, and the mean classification accuracy across all iterations was used as the final performance estimate for the model in each dataset.

Within each cross-validation iteration, the training folds were used to fit the global isoform- and gene-based RF models, while the held-out fold served as the independent test set for evaluation. The test cells were first used to assess global model performance and to identify concordant (overlapping) and discordant (non-overlapping) predictions between isoform- and gene-based classifiers. The same training folds were then used to construct subset-specific models, which were subsequently evaluated on the non-overlapping test cells. This hierarchical evaluation framework enabled systematic assessment of model performance at both global and subset levels within each cross-validation split (Figure 1).

**Figure 1:**
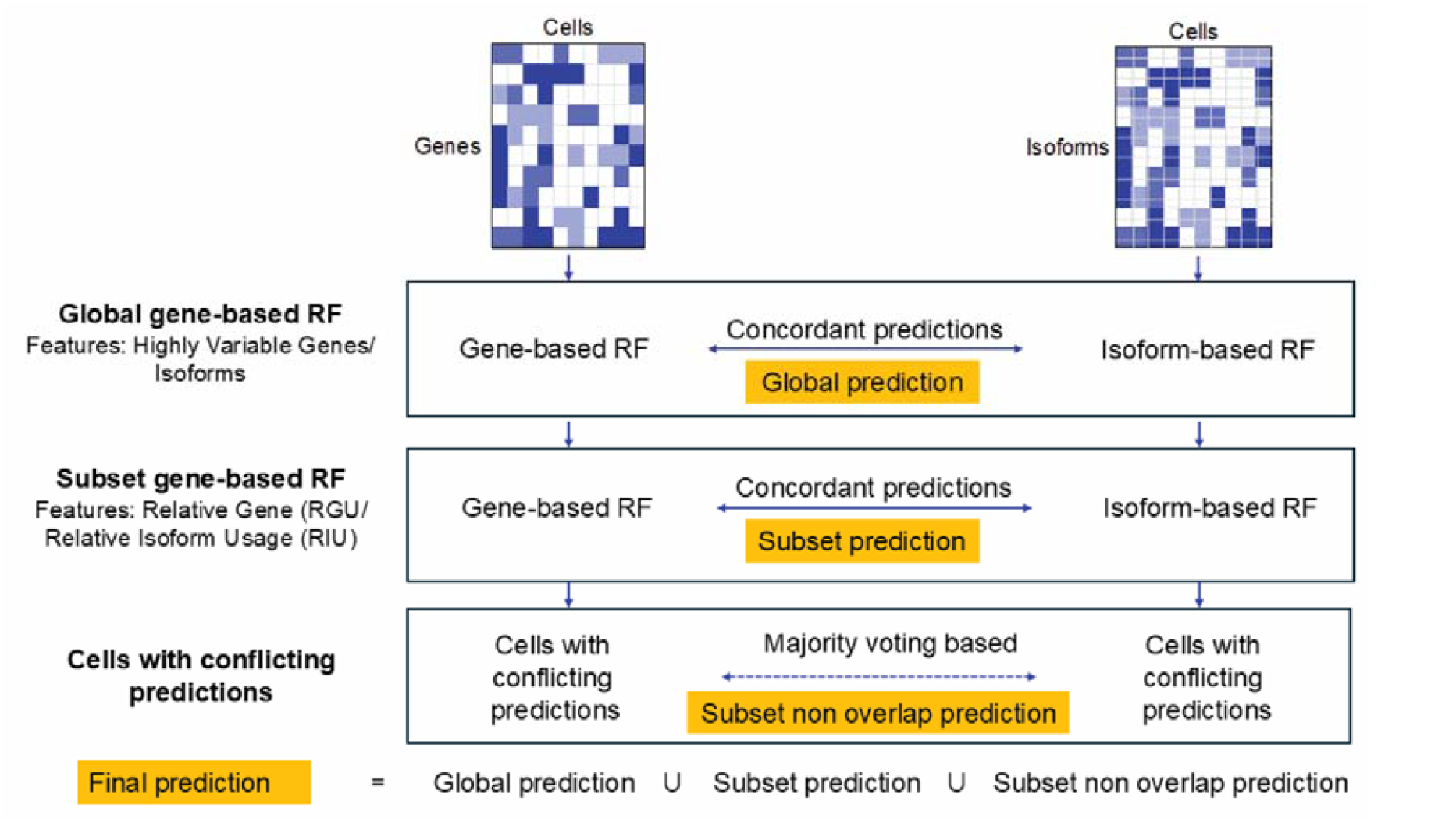
Feature engineering–based overlapping hierarchical random forest strategy.

#### Global models

At each iteration, four folds were used for training and the remaining fold was used for testing. Feature selection was performed independently for each model: highly variable isoforms were used for the isoform-based RF model, and highly variable genes were used for the gene-based RF model. Predictions were generated on the held-out test fold, and cells for which both models assigned the same cell type were defined as global overlapping predictions and retained as high-confidence assignments.

#### Subset models

Subset models were developed to refine predictions specifically for the non-overlapping cells produced by the global models. These models were trained using the same four training folds used for the global models, but with a different feature selection strategy: top relative isoform usage (RIU) for the isoform-based model and top relative gene usage (RGU) for the gene-based model. The subset models were then tested exclusively on the non-overlapping global test cells. For cells where both subset models agreed, the shared label was taken as the final prediction; for remaining disagreements, only the isoform-based label was retained due to its higher discriminative power at isoform resolution.

Based on this strategy, each test cell is assigned a prediction at only one level: global overlap, subset overlap, or subset non-overlap. Cells that do not have overlapping predictions at the global level proceed to the subset levels; similarly, only cells without overlapping predictions at the subset level are considered for the subset non-overlap predictions. The final prediction for each cell is therefore taken directly from the level at which it is classified (global, subset, or subset non-overlap), ensuring that every cell receives a single, unique prediction derived from the hierarchical strategy.

### 2.6 Benchmark of hierarchical random forest compared to gene-based annotation tools

To ensure a fair comparison, benchmarking was conducted separately using gene-level and isoform-level count matrices. In each setting, the selected annotation tools (Table 1) were trained independently on either long-read–derived gene-level matrices or isoform-level matrices and evaluated on corresponding held-out test datasets. Identical preprocessing, training, and evaluation procedures were applied in both cases to maintain methodological consistency. This enabled a direct comparison of existing cell type annotation tools across gene-based and isoform-based datasets.

**Table 1:** Cell type annotation tools included in the benchmark study. Internal: preprocessing is performed during tool execution; External: data must be preprocessed prior to input into the tool. The ‘Unassigned’ function represents whether the tool assigns cells that are uncertain in classification, such as those not meeting the predefined threshold of the respective tool.

| Tool | Classifier / Method | Predicts New Cell Types | Unassigned Function | Input Format | Preprocessing | Language | Reference |
| --- | --- | --- | --- | --- | --- | --- | --- |
| *SingleCellNet* | Random Forest | Yes | Yes | Expression matrices + cell metadata | External | R, Python | [22] |
| *scPred* | Support Vector Machine | No | Yes | *Seurat* object | Internal | R | [23] |
| *SingleR* | Spearman correlation (cell-to-reference) | No | No | SingleCellExperiment object | External | R | [24] |
| *clustifyR* | Spearman correlation (cluster-to-reference) | Yes | Yes | Reference: *Seurat* object +<br>Query: Clustered *Seurat* object | Internal | R | [25] |

#### Training and preprocessing

For all datasets, 70% of cells were used for training and 30% for testing. Each tool’s input was pre-processed according to the tool’s requirements (Table 1).

##### Bootstrapping for the early blood development and brain datasets

As both datasets contain fewer than 500 cells, several of the annotation tools benchmarked in this study showed reduced predictive performance when trained on such limited sample sizes. To mitigate this issue, bootstrapping was applied to the training data, while the test data were kept unchanged to ensure unbiased evaluation.

For both the datasets, 40% of cells from each cell type were randomly selected for training, and the remaining 60% were reserved for testing. To balance the training set, cell types with fewer than 120 cells were augmented by sampling additional cells with replacement to reach 150 cells. Duplicate cells were assigned unique barcodes, producing a final training set containing both original and synthetic cells. This ensured balanced representation of all cell types while maintaining unique identifiers for downstream analysis.

##### Clustering of Query Data for clustifyR

As *clustifyR* requires a pre-clustered dataset to assign cell types at the cluster level, the standard *seurat* clustering workflow was applied to each dataset to generate the clusters for the query data.

#### Benchmarking procedure

Each tool was trained separately on gene-level and isoform-level count matrices and evaluated on the corresponding held-out test cells for each dataset. Accuracy was calculated as the proportion of correctly classified cells across the test sets for each data representation.

## 3. Results

We developed an isoform-based HRF framework, ICON (Isoform-aware Cell classificatiON), for cell type annotation by incorporating feature engineering at both gene and isoform levels. A key feature introduced in this study is relative isoform usage (RIU), which quantifies the contribution of each isoform to a given cell type by measuring its expression proportion relative to the total isoform expression within that cell type. In parallel, relative gene usage (RGU) was computed to capture the proportional contribution of each gene within a cell type, providing a complementary gene-level representation.

The proposed model operates in a two-stage hierarchical manner. In the first stage, two independent RF classifiers are trained in parallel at the global level using gene and isoform count matrices, respectively, with features selected from highly variable genes and isoforms. Initial cell type predictions are generated independently by both models. Cells are then assigned to subset-level models based on an overlapping consensus strategy derived from agreement between gene- and isoform-level predictions, while non-overlapping (discordant) cells are flagged for further refinement (Figure 1).

In the second stage, these discordant cells are re-evaluated using RIU- and RGU-based representations, which more sensitively capture cell type–specific expression patterns (Figure 1). For these non-overlapping cells, predictions are refined using majority voting based on isoform-level models. Final cell type assignments are obtained by integrating predictions across all three levels of the framework (global, subset, and non-overlapping refinement stages), where each test cell receives a prediction from the appropriate stage depending on its pathway through the hierarchy (Figure 1). This hierarchical and integrative framework enables more robust assignment of cells to subtype-specific models and improves overall annotation accuracy.

### 3.1 Informative features have greater impact on RF performance than increased model complexity

To optimise the performance of RF classifiers for cell type annotation, we systematically evaluated the impact of key model parameters, focusing primarily on the number of trees and the number of input features. Using the LongBench PacBio and Nanopore isoform count matrices, which provide a sufficiently large number of cells for robust training and testing, we assessed how varying these parameters influenced classification accuracy, macro F1 score, and weighted F1 score. This analysis enabled identification of parameter settings that balance predictive performance and computational efficiency, providing a foundation for subsequent model development.

Increasing the number of trees from 100 to 1,000 led to a visible improvement in the macro F1 score. However, the other two evaluation metrics, classification accuracy and weighted F1 score, varied by less than 1% across this range (Figure 2A-C). Because macro F1 assigns equal weight to all classes and does not account for class imbalance, it is particularly sensitive to rare cell types. In contrast, weighted F1 reflects overall performance in imbalanced single-cell datasets by weighting classes according to their prevalence. Therefore, for the selection of optimal number of features and trees, we prioritised weighted F1 score and classification accuracy.

**Figure 2:**
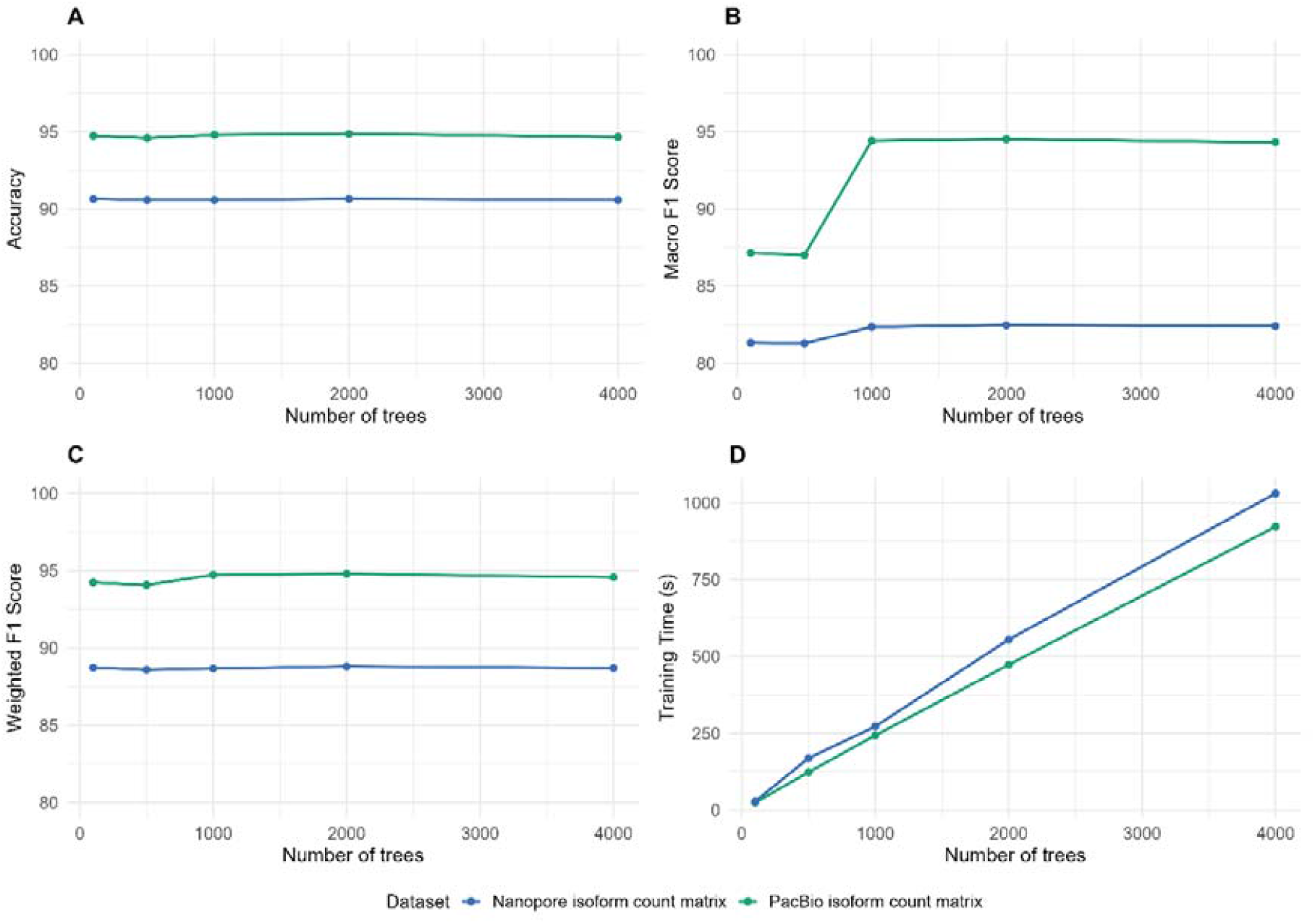
Effect of the number of trees on random forest performance with a fixed feature set. Panels show changes in **(A)** accuracy, **(B)** macro F1 score, **(C)** weighted F1 score, and **(D)** training time as the number of trees increases. These trends were used to determine an appropriate number of trees that balances predictive performance and computational cost.

Given the substantial increase in training time associated with larger numbers of trees and the marginal gains observed in performance metrics, we selected 100 trees as a computationally efficient and robust configuration for all subsequent analyses.

Model performance varied depending on the number of features used (Figure 3). Based on these results, 1,000 features were selected for RF training, as this configuration achieved strong classification performance without introducing unnecessary model complexity. Increasing the number of features beyond 1,000 led to only marginal improvements in accuracy and weighted F1 score (<1%), while substantially increasing computational cost. These results emphasise that the informativeness of features, rather than their quantity, drives classification performance.

**Figure 3:**
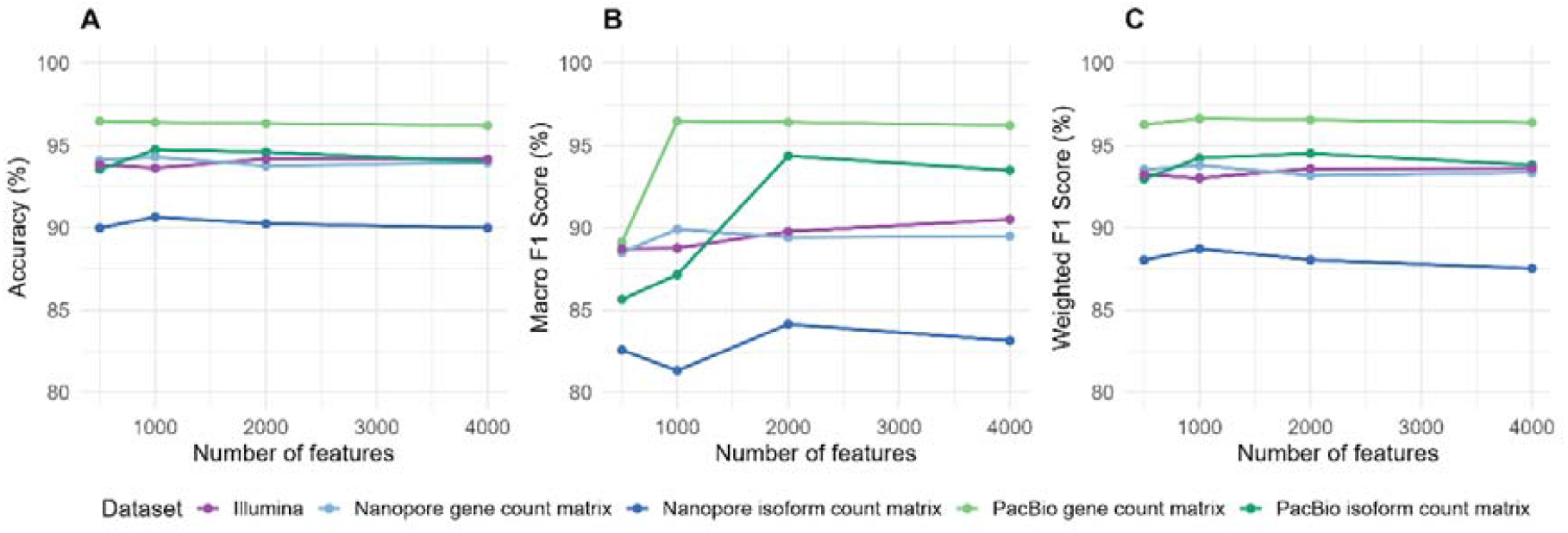

### 3.2 Hierarchical integration of isoform and gene features improves cell type annotation performance

ICON integrates isoform- and gene-level information for cell type annotation through a multi-stage classification strategy combining global and subset models. The results below summarise the performance of ICON evaluated using a 5-fold cross-validation strategy across all datasets.

Across most datasets, the combined HRF predictions perform better than the isoform-based RF model alone (Figure 4). Interestingly, the combined strategy (final prediction) does not surpass the performance of the gene-based RF model. These results indicate that while the hierarchical approach improves upon isoform-only modelling, gene-level features continue to provide the strongest single-source predictive signal for these datasets.

**Figure 4:**
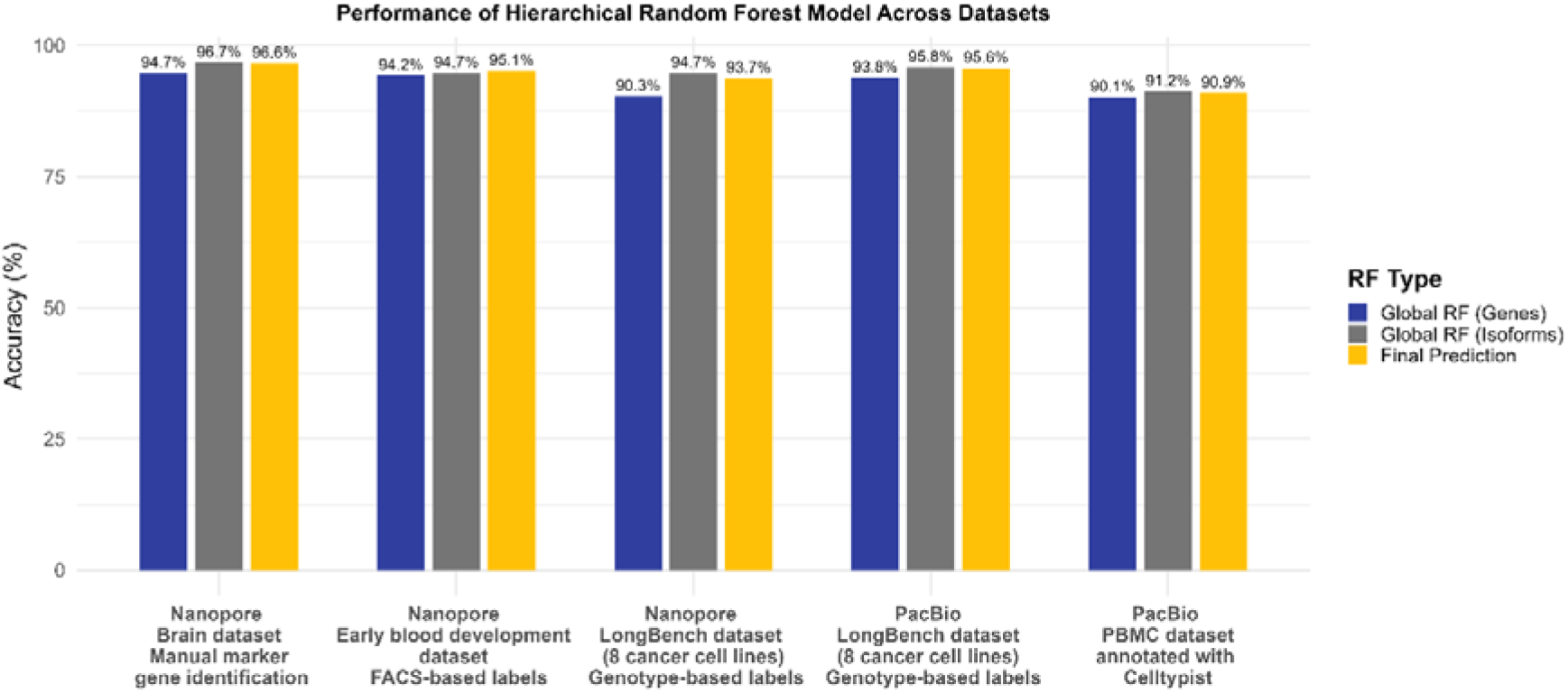
Performance of the hierarchical random forest model across datasets. The global-level random forest model results show how the model performs when using either isoform-based (grey) or gene-based (blue) count matrices independently. The yellow bar represents the final prediction accuracy of the hierarchical random forest model, which integrates both gene and isoform features.

### 3.3 Hierarchical integration of gene and isoform features surpasses conventional gene-based annotation tools

To evaluate the effectiveness of existing cell type annotation approaches on long-read scRNA-seq data, we benchmarked ICON against representative gene-based annotation tools using both gene-level and isoform-level count matrices across multiple long-read datasets with experimentally validated cell type labels. For each dataset, the tools were trained and evaluated separately on gene-based and isoform-based representations, enabling a direct comparison of performance across data modalities. These benchmarks were designed to assess how well conventional gene-level methods generalise to isoform-level count matrices and to characterise performance differences attributable to data representation and tool design.

The benchmarking results demonstrate that the HRF consistently outperforms traditional gene-based annotation tools when applied to both gene-level and isoform-level count matrices (Figure 5A,B). Although gene-based methods could be retrained on isoform-level data, their performance was substantially reduced compared to their performance on gene-level matrices and remained lower than that of the HRF across datasets. This performance gap was consistently observed in all datasets, indicating limitations in directly adapting gene-centric models to isoform-level representations.

**Figure 5:**
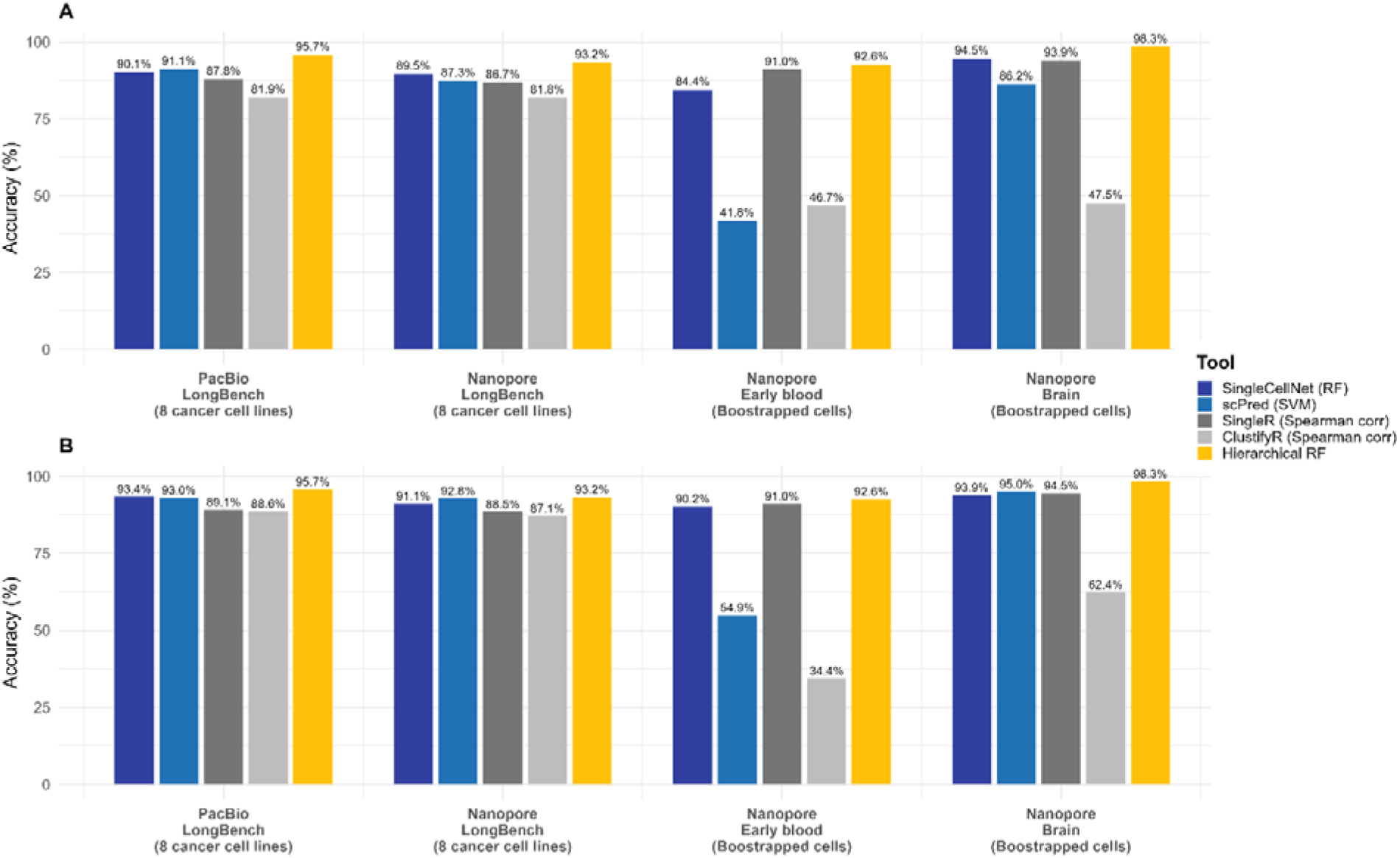
Benchmarking performance of cell type annotation tools across multiple datasets. **(A)** Results for isoform count matrix–based datasets. **(B)** Results for gene count matrix–based datasets. Bars represent classification accuracy, calculated by comparing predicted cell type labels against experimentally validated ground-truth annotations.

Importantly, ICON was explicitly designed to incorporate isoform-based features while preserving gene-level structure and hierarchical relationships among cell types. Its superior performance when benchmarked across multiple gene- and isoform-level datasets demonstrates that integrating gene- and isoform-level information substantially improves the accuracy of cell type annotation in long-read scRNA-seq data.

### 3.4 Key predictive features revealed through model interpretability uncover black-box architecture

To identify the features (isoforms and genes) underlying cell type discrimination, RF variable importance scores were extracted from the global and subset-level HRF models, and isoforms/genes were ranked within each cell type according to class-specific importance scores, where higher scores indicate greater contribution to accurate prediction of the corresponding cell type. This enabled direct comparison of the most discriminative features across isoform- and gene-level representations in all four models (global isoform, global gene, subset isoform, and subset gene). The ranked feature profiles revealed both shared and model-specific predictive signatures across classification stages.

These highly informative features may also represent candidate biomarkers for future cell type annotation following further biological and experimental validation. Cell type–specific important features derived from the PacBio PBMC dataset for four cell types are shown in Figure 6, with results for all datasets provided in the Supplementary Materials.

**Figure 6:**
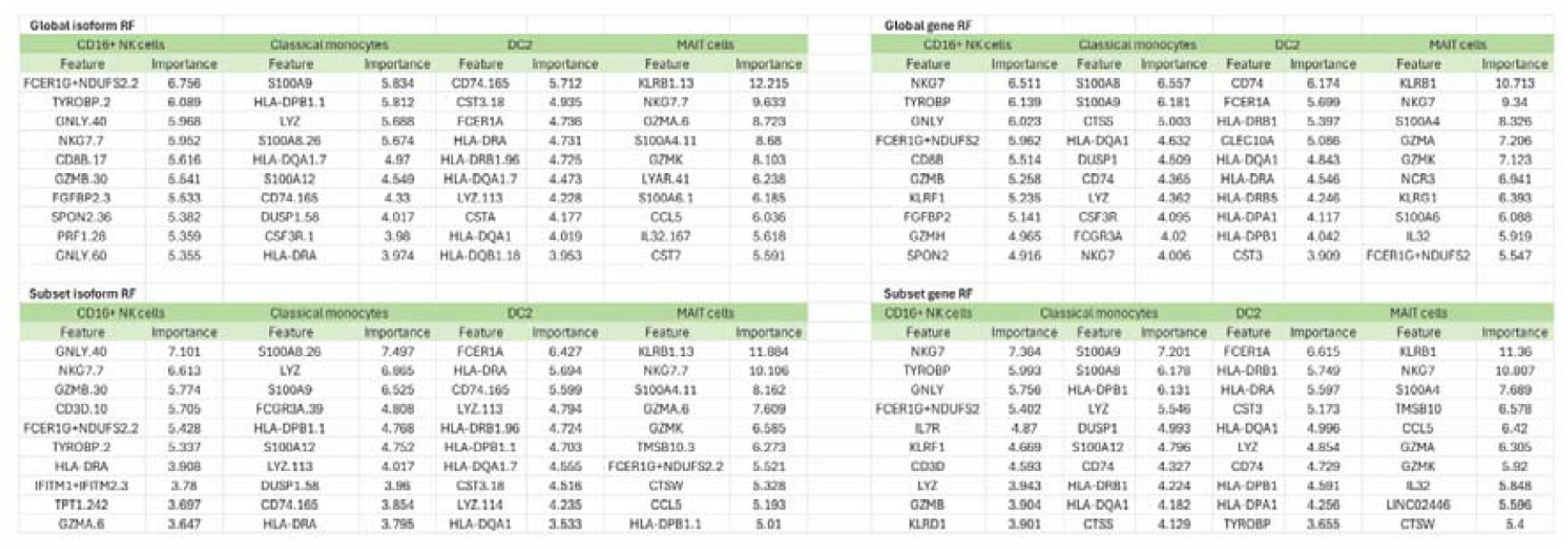
Cell type–specific important features identified from the PacBio PBMC dataset. The figure shows the most informative features contributing to the classification of four representative PBMC cell types, highlighting genes and isoforms with high importance scores derived from the model.

## 4. Discussion

### 4.1 Model development and performance evaluation

We developed a HRF model (ICON) for cell type annotation using isoform-level information. The feature engineering–based HRF framework enables interpretation of model predictions, with particular emphasis on the contributions of gene- and isoform-level features.

Due to the current lack of cell type annotation tools specifically designed for long-read scRNA-seq data, particularly for isoform-based count matrices, this study focused on developing and evaluating a methodology tailored to isoform-level cell type classification. To contextualise the performance of ICON, it has been benchmarked against representative gene-based annotation tools (Table 1). Although both *singleR* [24] and *clustifyR* [25] use Spearman correlation for cell type classification, *clustifyR* was specifically chosen because, unlike the other three tools that provide predictions at the individual cell level, *clustifyR* generates cell type annotations at the cluster level.

Most existing cell type annotation tools are designed for short-read scRNA-seq data and are trained using gene-level expression matrices [15]. However, long-read sequencing technologies provide transcript- and isoform-level resolution, generating isoform count matrices in which feature identifiers no longer correspond directly to gene names. As a result, pre-trained gene-based annotation models cannot be directly applied to isoform-level data, and the suitability of existing tools for this data modality remains unclear.

Overall, our benchmarking framework exposed the limitations of traditional gene-based approaches when applied to isoform-level data and highlighted the need for annotation strategies that explicitly leverage isoform-level information.

Feature engineering based HRF model incorporated a combined gene- and isoform-based overlapping prediction strategy. The introduction of novel engineered features, relative isoform usage and relative gene usage, enabled subset models to be fine-tuned in a more cell type specific manner. These features captured cell type dependent expression patterns more effectively and improved discrimination between closely related cell types.

Although the feature engineering–based HRF model did not consistently outperform gene-only RF models across all datasets, it demonstrated higher performance than existing gene-based cell type annotation tools when applied to both isoform-level and gene-level count matrices (based on our benchmark). The results also indicate that gene-based tools generally perform well on gene-level data, achieving high accuracy for most cell types. However, their performance drops substantially on isoform-level matrices, highlighting a key limitation of gene-centric methods. While these tools are effective for short-read gene expression data, their underlying assumptions and feature representations do not translate well to long-read isoform-based data. Simply retraining gene-based models on isoform identifiers does not adequately capture the biological and structural complexity introduced by transcript-level resolution, resulting in reduced classification accuracy.

The consistently improved performance of the HRF underscores the importance of developing annotation methods specifically tailored to long-read sequencing data. Isoform-level expression contains rich, cell type specific information that is not fully represented at the gene level, and effective utilisation of this information requires models designed with isoform-based features in mind [9,26]. Overall, these findings demonstrate the need for dedicated cell type annotation tools for long-read, isoform-based scRNA-seq data. ICON provides a step toward addressing this gap and establishes a foundation for future method development in isoform-aware cell type annotation

### 4.2 Interpretation of the hierarchical random forest model

ICON improves model interpretability by enabling feature-level interpretation across both global and subset classification stages. Unlike many black-box ML approaches, which make predictions without providing clear insight into how features influence the output [27], RF-based variable importance scores provide direct insight into the features driving cell type discrimination, with higher importance values reflecting stronger predictive influence for specific cell types. In this study, the incorporation of relative isoform and gene usage further enhanced interpretability by capturing cell type–specific expression patterns at a finer resolution, enabling more biologically meaningful feature prioritisation within subset models. Importantly, the hierarchical structure allows comparison of feature contributions across isoform- and gene-level models trained on identical biological samples, facilitating the identification of consistent and discordant molecular signatures. These prioritised isoforms and genes may provide promising marker candidates for future cell type annotation, pending additional biological and experimental validation. This multi-level interpretability is particularly valuable for long-read RNA sequencing data, where isoform resolution reveals regulatory complexity that is not observable at the gene level alone.

## Authors contributions

HW surveyed the literature and wrote the first draft. SW and US supervised the work and helped with reviewing and revising the manuscript. All authors read and approved the final manuscript.

## Code availability

The R-based workflow used in this study for cell type annotation in single-cell long-read RNA-seq data, implementing the feature engineering–based HRF framework with isoform- and gene-level predictions, is publicly available at https://github.com/jithmaWijeward1/

## References

[1] Reyes A, Huber W. Alternative start and termination sites of transcription drive most transcript isoform differences across human tissues. Nucleic Acids Res 2018;46:582–92. 10.1093/nar/gkx1165.

[2] Wang H, Yang Y, Liu J, Qian L. Direct cell reprogramming: approaches, mechanisms and progress. Nat Rev Mol Cell Biol 2021;22:410–24. 10.1038/s41580-021-00335-z.

[3] McIntire E, Barr KA, Gonzales NM, Gilad Y. Guided Differentiation of Pluripotent Stem Cells for Cardiac Cell Diversity. BioRxiv Preprint 2025. 10.1101/2023.07.21.550072.

[4] Sadelain M, Rivière I, Riddell S. Therapeutic T cell engineering. Nature 2017;545:423–31. 10.1038/nature22395.

[5] Lebrigand K, Magnone V, Barbry P, Waldmann R. High throughput error corrected Nanopore single cell transcriptome sequencing. Nat Commun 2020;11:4025. 10.1038/s41467-020-17800-6.

[6] Gupta P, O’neill H, Wolvetang EJ, Chatterjee A, Gupta I. Advances in single-cell long-read sequencing technologies. NAR Genom Bioinform 2024;6. 10.1093/nargab/lqae047.

[7] Stark R, Grzelak M, Hadfield J. RNA sequencing: the teenage years. Nat Rev Genet 2019;20:631–56. 10.1038/s41576-019-0150-2.

[8] Mincarelli L, Uzun V, Wright D, Scoones A, Rushworth SA, Haerty W, et al. Single-cell gene and isoform expression analysis reveals signatures of ageing in haematopoietic stem and progenitor cells. Commun Biol 2023;6:558. 10.1038/s42003-023-04936-6.

[9] Gupta I, Collier PG, Haase B, Mahfouz A, Joglekar A, Floyd T, et al. Single-cell isoform RNA sequencing characterizes isoforms in thousands of cerebellar cells. Nat Biotechnol 2018;36:1197–202. 10.1038/nbt.4259.

[10] Monzó C, Liu T, Conesa A. Transcriptomics in the era of long-read sequencing. Nat Rev Genet 2025;26:681–701. 10.1038/s41576-025-00828-z.

[11] Bhatia S, Field MA, Hebbard L, Schmitz U. Bioinformatics frameworks for single-cell long-read sequencing: unlocking isoform-level resolution. Brief Bioinform 2025;26. 10.1093/bib/bbaf655.

[12] Wu S, Schmitz U. ScIsoX: a multidimensional framework for measuring isoform-level transcriptomic complexity in single cells. Genome Biol 2025;26:289. 10.1186/s13059-025-03758-5.

[13] Wu S, Schmitz U. Single-cell and long-read sequencing to enhance modelling of splicing and cell-fate determination. Comput Struct Biotechnol J 2023;21:2373–80. 10.1016/j.csbj.2023.03.023.

[14] Method of the Year 2022: long-read sequencing. Nat Methods 2023;20:1. 10.1038/s41592-022-01759-x.

[15] Wijewardena H, Bhatia S, Bhattacharya N, Sengupta D, Wu S, Schmitz U. Advancing automated cell type annotation with large language models and single-cell isoform sequencing. Comput Struct Biotechnol J 2025;27:4952–62. 10.1016/j.csbj.2025.11.008.

[16] Clarke ZA, Andrews TS, Atif J, Pouyabahar D, Innes BT, MacParland SA, et al. Tutorial: guidelines for annotating single-cell transcriptomic maps using automated and manual methods. Nat Protoc 2021;16:2749–64. 10.1038/s41596-021-00534-0.

[17] Chen J, Zhang J, Yao H, Li Y. CellTypeAgent: Trustworthy cell type annotation with Large Language Models. ArXiv Preprint 2025. 10.48550/arXiv.2505.08844.

[18] Satija R, Farrell JA, Gennert D, Schier AF, Regev A. Spatial reconstruction of single-cell gene expression data. Nat Biotechnol 2015;33:495–502. 10.1038/nbt.3192.

[19] Sant C, Mucke L, Corces MR. CHOIR improves significance-based detection of cell types and states from single-cell data. Nat Genet 2025. 10.1038/s41588-025-02148-8.

[20] Ma F, Pellegrini M. ACTINN: Automated identification of cell types in single cell RNA sequencing. Bioinformatics 2020;36:533–8. 10.1093/bioinformatics/btz592.

[21] Lieberman Y, Rokach L, Shay T. CaSTLe - Classification of single cells by transfer learning: Harnessing the power of publicly available single cell RNA sequencing experiments to annotate new experiments. PLoS One 2018;13:e0205499. 10.1371/journal.pone.0205499.

[22] Tan Y, Cahan P. SingleCellNet: A Computational Tool to Classify Single Cell RNA-Seq Data Across Platforms and Across Species. Cell Syst 2019;9:207-213.e2. 10.1016/j.cels.2019.06.004.

[23] Alquicira-Hernandez J, Sathe A, Ji HP, Nguyen Q, Powell JE. ScPred: Accurate supervised method for cell-type classification from single-cell RNA-seq data. Genome Biol 2019;20:264. 10.1186/s13059-019-1862-5.

[24] Aran D, Looney AP, Liu L, Wu E, Fong V, Hsu A, et al. Reference-based analysis of lung single-cell sequencing reveals a transitional profibrotic macrophage. Nat Immunol 2019;20:163–72. 10.1038/s41590-018-0276-y.

[25] Fu R, Gillen AE, Sheridan RM, Tian C, Daya M, Hao Y, et al. clustifyr: an R package for automated single-cell RNA sequencing cluster classification. F1000Res 2020;9:223. 10.12688/f1000research.22969.1.

[26] Tilgner H, Jahanbani F, Blauwkamp T, Moshrefi A, Jaeger E, Chen F, et al. Comprehensive transcriptome analysis using synthetic long-read sequencing reveals molecular co-association of distant splicing events. Nat Biotechnol 2015;33:736–42. 10.1038/nbt.3242.

[27] Dunn J, Mingardi L, Zhuo YD. Comparing interpretability and explainability for feature selection. ArXiv Preprint 2021. 10.48550/arXiv.2105.05328.

