## Supplementary Materials for "ICON: An isoform-aware hierarchical random forest model for cell type classification"

Supplementary Table 1: Datasets utilised for tool development and evaluation.

| **Dataset** | **Description** | **Cell Type Annotation** | **Cell Count** | **Technology** | **Reference** |
| --- | --- | --- | --- | --- | --- |
| Brain dataset | Major neuronal and non-neuronal brain cell types | Manual marker gene identification | 1231 (bootstrapped) | ONT | [18] |
| Early Blood Development | Murine hematopoietic development | Experimentally validated FACS-isolated embryonic cell labels | 1172  (bootstrapped) | ONT | [19] |
| LongBench | 8 cancer cell lines | Genotype-based cell line annotations | ONT: 5023, PacBio: 5002 | ONT, PacBio, Illumina | [20] |
| PacBio PBMC | Peripheral blood immune cell populations | *Celltypist* based cell type annotation | 8816 | PacBio | [21] |

ONT – Oxford Nanopore Technologies; PacBio – Pacific Biosciences; FACS - Fluorescence-activated cell sorting; PBMC - peripheral blood mononuclear cells.

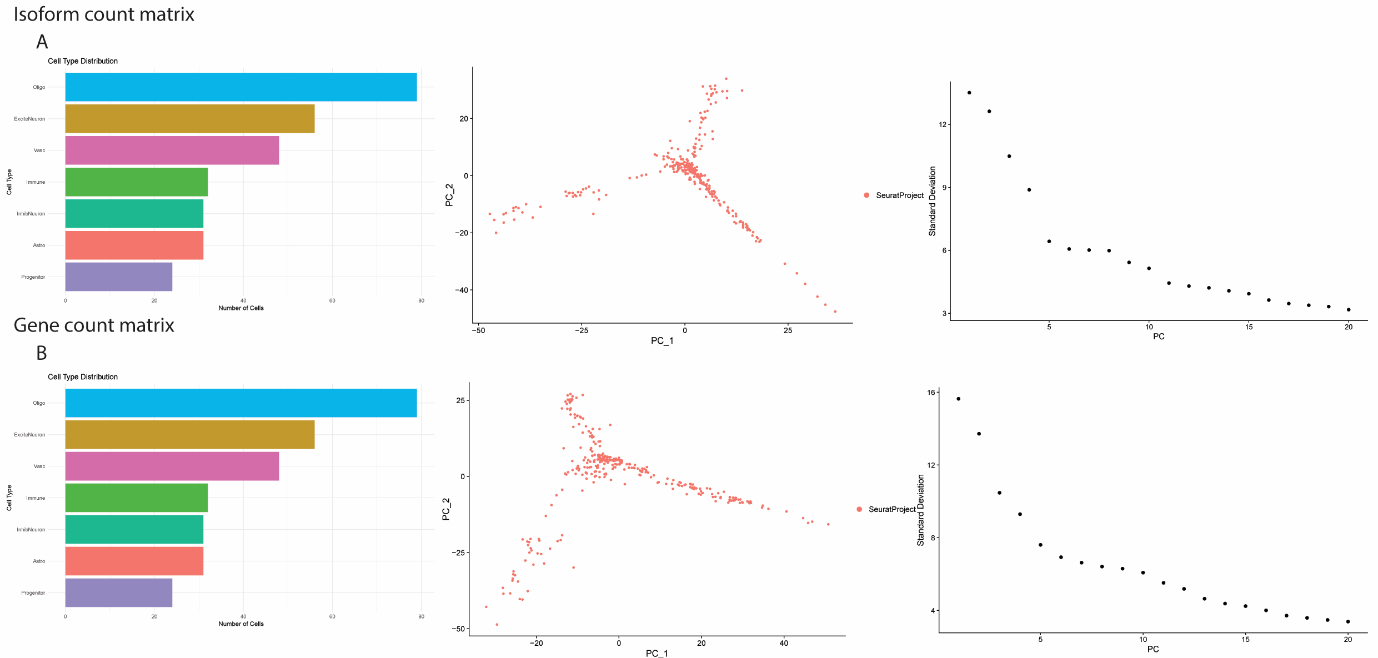

Figure 1: Brain dataset comparative overview of isoform and gene expression profiles across single-cell RNA-seq data. (A) Isoform count matrix analysis, including cell type distribution (bar chart), dimensionality reduction using PCA, and elbow plot for optimal component selection. (B) Gene count matrix analysis, showing corresponding cell type distribution (bar chart), PCA-based visualization, and elbow plot for dimensionality assessment.

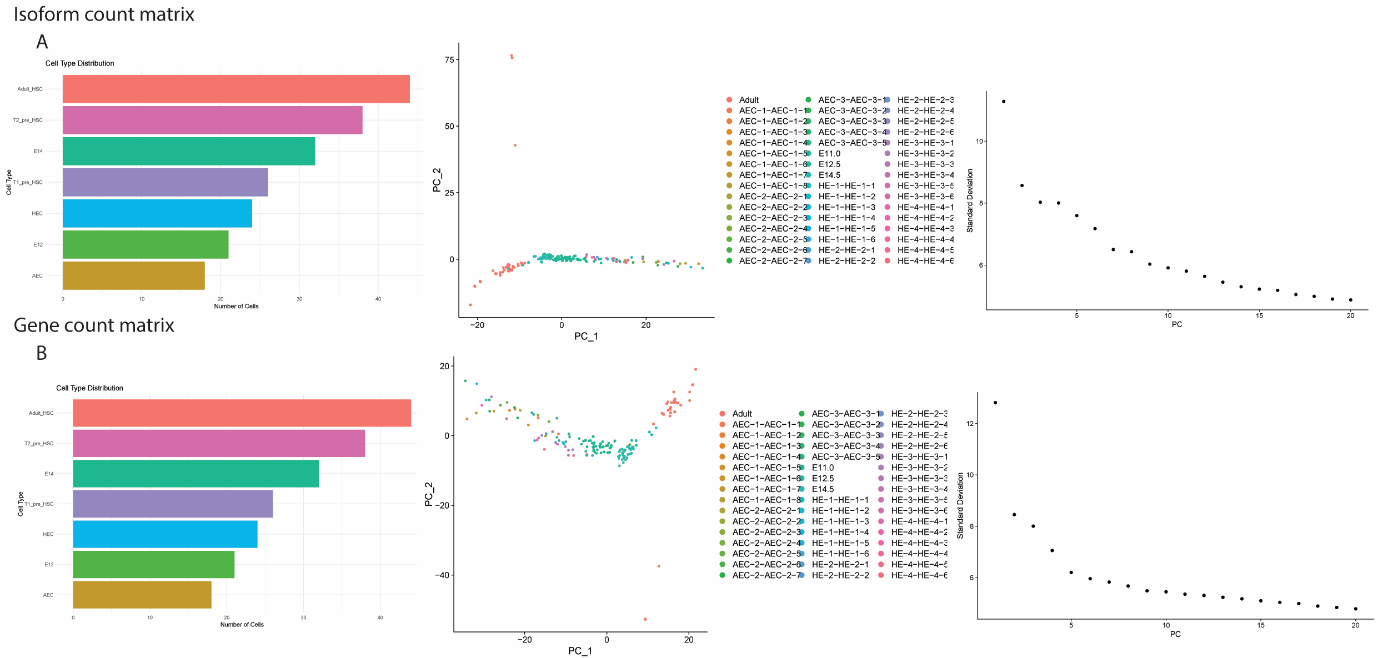

Figure 2: Early blood dataset comparative overview of isoform and gene expression profiles across single-cell RNA-seq data. (A) Isoform count matrix analysis, including cell type distribution (bar chart), dimensionality reduction using PCA, and elbow plot for optimal component selection. (B) Gene count matrix analysis, showing corresponding cell type distribution (bar chart), PCA-based visualization, and elbow plot for dimensionality assessment.

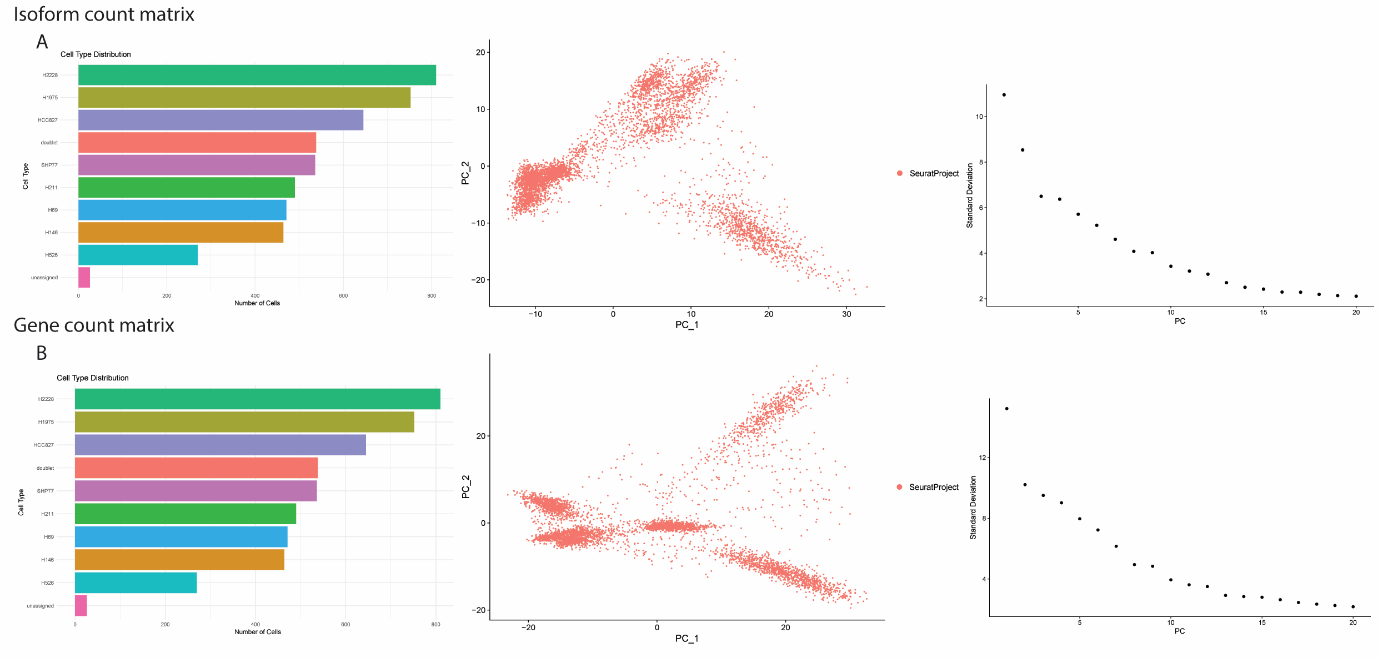

Figure 3: LongBench PacBio dataset comparative overview of isoform and gene expression profiles across single-cell RNA-seq data. (A) Isoform count matrix analysis, including cell type distribution (bar chart), dimensionality reduction using PCA, and elbow plot for optimal component selection. (B) Gene count matrix analysis, showing corresponding cell type distribution (bar chart), PCA-based visualization, and elbow plot for dimensionality assessment.

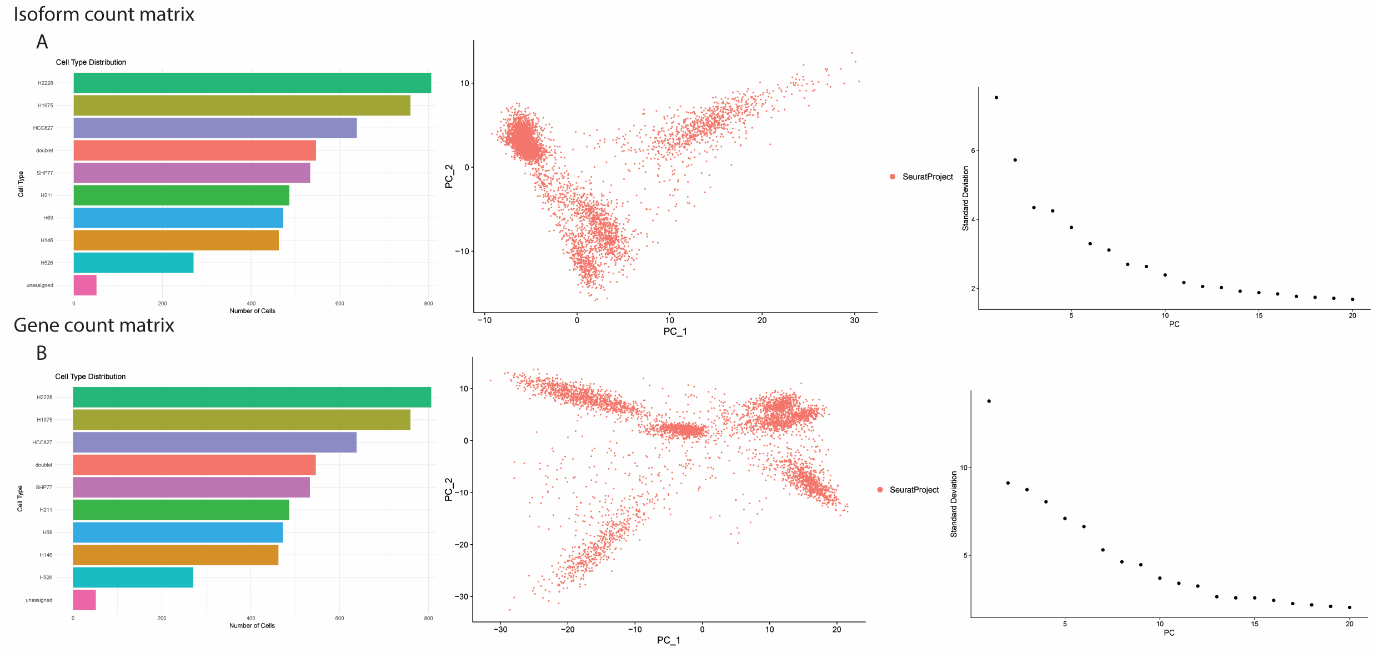

Figure 4: LongBench Nanopore dataset comparative overview of isoform and gene expression profiles across single-cell RNA-seq data. (A) Isoform count matrix analysis, including cell type distribution (bar chart), dimensionality reduction using PCA, and elbow plot for optimal component selection. (B) Gene count matrix analysis, showing corresponding cell type distribution (bar chart), PCA-based visualization, and elbow plot for dimensionality assessment.

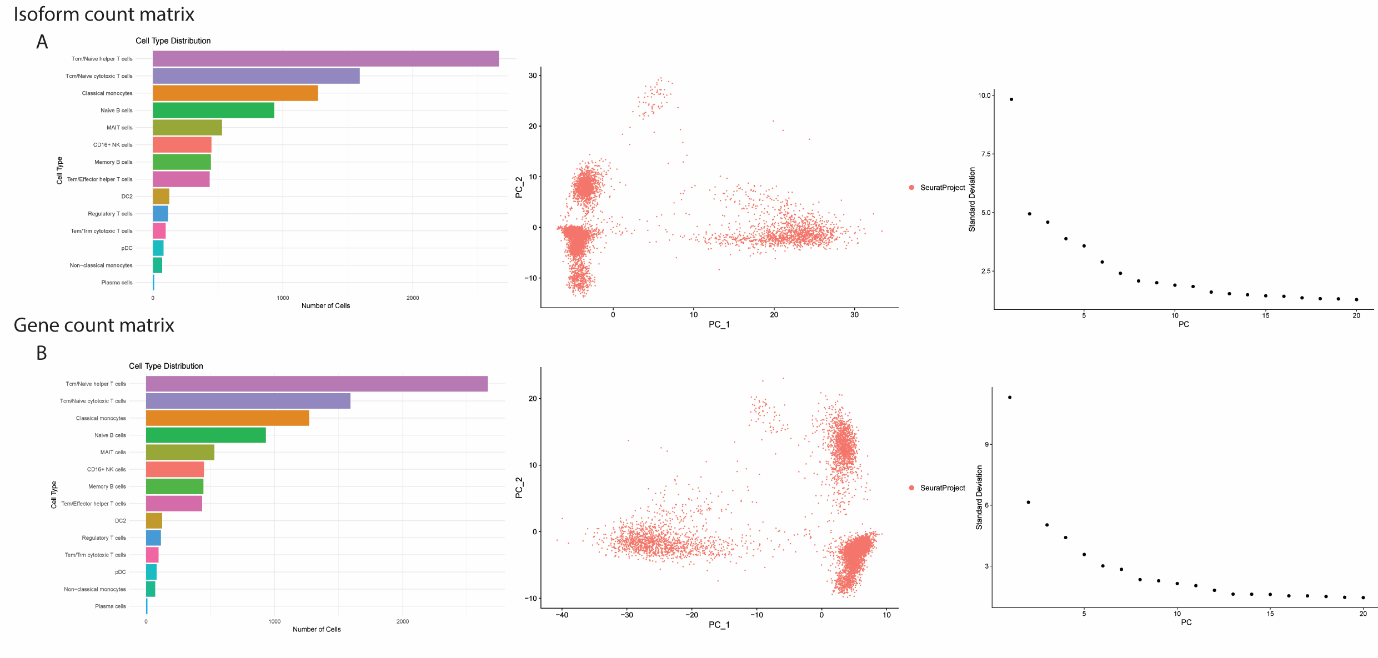

Figure 5: PacBio PBMC dataset comparative overview of isoform and gene expression profiles across single-cell RNA-seq data. (A) Isoform count matrix analysis, including cell type distribution (bar chart), dimensionality reduction using PCA, and elbow plot for optimal component selection. (B) Gene count matrix analysis, showing corresponding cell type distribution (bar chart), PCA-based visualization, and elbow plot for dimensionality assessment.

### **Extended Methods**

#### **Random Forest model parameter optimisation**

Before implementing the HRF model we evaluated RF hyperparameters using LongBench PacBio and Nanopore isoform count matrices (Supplementary table 1). These datasets were selected because, compared with other experimentally validated cell-type–annotated datasets we use in this study, these datasets contain a relatively large number of cells (>5,000), providing sufficient samples for robust model training and testing.

To determine an appropriate number of trees, we trained RF models with varying tree counts (100, 500, 1’000, 2’000, and 4’000), while fixing the number of input features (isoforms) at 1’000. Model performance was evaluated using classification accuracy, macro F1 score, and weighted F1 score.

Both isoform- and gene-level count matrices contain a very large number of features, of which only a subset is informative for cell type annotation. Consequently, the choice of the number of input features can strongly influence model performance. To investigate this effect, we varied the number of features (500, 1’000, 2’000, and 4’000) across the LongBench, PacBio, and Nanopore datasets, using both isoform and gene-level count matrices. Model performance was assessed using classification accuracy, macro F1 score, and weighted F1 score.

###### **Relative isoform usage and relative gene usage**

To support the hierarchical feature engineering framework and to identify the most informative isoforms and genes for cell type classification, we introduced two novel feature representations: relative isoform usage (RIU) and relative gene usage (RGU). These values indicate how strongly a specific isoform or gene is expressed within a given cell type, serving as features that help distinguish one cell type from another. To calculate these values, cells were first stratified by cell type, and for each cell type, the relative isoform or gene usage was computed as the proportion of an isoform’s or gene’s expression relative to the total expression of all isoforms or genes in that cell type.

Let *i* denote the specific isoform of interest, *j* an index variable used for summation over all isoforms, and *J* the total number of isoforms considered within a given cell type *c*. The relative isoform usage for isoform *i* in cell type *c* was defined as:

$$\text{Relative Isoform Usage}_{\text{i,c}}=\frac{\text{Expression of isoform }i\text{ in cell type }c}{\sum_{j=1}^{J} \text{Expression of isoform }j\text{ in cell type }c}$$

The relative gene usage was calculated in a similar manner, by taking the proportion of a gene’s expression relative to the total expression of all genes in the same cell type.

Isoforms and genes were then ranked in descending order based on their relative isoform and gene usage value within each cell type. The top-ranking isoforms for each cell type, representing cell type–specific isoform usage, were selected and then combined into a single set of candidate isoforms. To ensure uniqueness, duplicate isoforms that appeared in multiple cell types were retained only once in the final list of top RIUs and RGUs.
